# Affected cell types for hundreds of Mendelian diseases revealed by analysis of human and mouse single-cell data

**DOI:** 10.1101/2022.10.29.513906

**Authors:** Idan Hekselman, Assaf Vital, Maya Ziv-Agam, Lior Kerber, Esti Yeger-Lotem

**Affiliations:** Department of Clinical Biochemistry and Pharmacology, Faculty of Health Sciences, Ben-Gurion University of the Negev, Be’er Sheva, Israel; The National Institute for Biotechnology in the Negev, Ben-Gurion University of the Negev, Be’er Sheva, Israel

**Keywords:** Single-cell transcriptomics, Hereditary diseases, Tissue and cell-type selectivity, Data integration

## Abstract

Hereditary diseases manifest clinically in certain tissues, however their affected cell types typically remain elusive. Single-cell expression studies showed that overexpression of disease-associated genes may point to the affected cell types. Here, we developed a method that infers disease-affected cell types from the preferential expression of disease-associated genes in cell types (PrEDiCT). We applied PrEDiCT to single-cell expression data of six human tissues, to infer the cell types affected in 1,113 hereditary diseases. Overall, we identified 110 cell types affected by 714 diseases. We corroborated our findings by literature text-mining and recapitulation in mouse corresponding tissues. Based on these findings, we explored features of disease-affected cell types and cell classes, highlighted cell types affected by mitochondrial diseases and heritable cancers, and identified diseases that perturb intercellular communication. This study expands our understanding of disease mechanisms and cellular vulnerability.

## INTRODUCTION

Hereditary diseases affect 6% of the world population and typically lack cure. Identification of their genetic and molecular basis is challenging, limiting their diagnosis and treatment (Ferreira, 2019). Knowledge of tissues and cell types that manifest with pathophysiological changes in patients was shown to facilitate the understanding of disease mechanisms (Hekselman & Yeger-Lotem, 2020). For example, transcriptomic analysis of muscle tissues helped to genetically diagnose patients with rare muscle disorders (Cummings et al., 2017). Likewise, identification of a rare cell type affected by cystic fibrosis opened new avenues for treatment (Montoro et al., 2018; Plasschaert et al., 2018). However, whereas affected tissues may be evident for many hereditary diseases, the exact cell types that manifest with pathophysiological changes within affected tissues are often unknown.

Owing to massive single-cell profiling of mammalian tissues (Tabula Sapiens, 2022), investigation of diseases in cellular contexts has become feasible. In particular, it was shown that genes whose aberration leads to disease (i.e., disease genes) tend to be expressed preferentially in cell types that express pathology (denoted disease-affected cell types). For instance, 21/29 mouse homologs of human disease genes associated with nephrotic syndrome were shown to be upregulated in podocytes, the disease-relevant renal cell type (Park et al., 2018). In an inspiring manner, preferential expression of the cystic fibrosis gene CFTR in ionocytes of human and mouse airways revealed their role in cystic fibrosis (Montoro et al., 2018; Plasschaert et al., 2018). Likewise, the preferential expression of disease genes for 18 Mendelian muscle disorders revealed disease-affected muscle cell types (Eraslan et al., 2022). Thus, preferential expression of disease genes may serve as an indicator of disease-affected cell types, and has the potential to shed light on the cellular mechanisms that underlie hereditary diseases.

Preferential expression of disease-associated genes has also been used to illuminate cell types affected by complex traits (Dai et al., 2021; Eraslan et al., 2022; Jagadeesh et al., 2021; Kim-Hellmuth et al., 2020). The CSEA-DB repository used enrichment analysis to associate complex traits to potentially-affected cell types (Dai et al., 2021). Other studies compared the differential expression of genes in cell types of diseased versus healthy tissues (Mathys et al., 2019; Segerstolpe et al., 2016; Smillie et al., 2019). SC2disease database associated 25 genetic diseases with hundreds of potentially-affected cell types via such comparisons (Zhao et al., 2021). However, most of these efforts rarely validated or corroborated the inferred associations.

Here, we developed a method that infers disease-affected cell types from the Preferential Expression of Disease genes in Cell Types (PrEDiCT). We applied PrEDiCT to 1,113 Mendelian diseases that manifest in six human tissues and inferred disease-affected cell types for 714 diseases. We corroborated our findings by text-mining of PubMed records and by their recapitulation in mice. The resulting large-scale resource allowed us to explore characteristics of disease-affected cell types across tissues, highlight cell types affected by mitochondrial diseases and heritable cancers, and identify diseases that perturb intercellular communication.

## RESULTS

### Preferential expression of disease genes indicates disease-affected cell types

To identify the cell types affected by Mendelian diseases with known disease genes, we developed the PrEDiCT scoring scheme (Fig. 1A). Below we describe the PrEDiCT workflow, including data acquisition, PrEDiCT score calculation, and validation. Data of annotated human single-cell transcriptomes were obtained from Tabula Sapiens (Tabula Sapiens, 2022). We focused on tissues with two or more samples with ≥800 sequenced cells that were also sequenced in mice [(Tabula Muris, 2018); Methods]. Tissues included bone marrow, lung, skeletal muscle, spleen, tongue and trachea, and altogether were comprised of 129 cell types. Next, we associated hereditary diseases to their affected tissues based on clinical records in Human Phenotype Ontology (HPO) database [(Kohler et al., 2021); Methods]. To verify HPO-based associations we compared them to ODiseA database, which contained manually curated associations between diseases and affected tissues [(Hekselman et al., 2022); Methods]. Indeed, most of HPO-based annotations were asserted by ODiseA (422 out of 548 associations tested, 77%). We then gathered the respective disease genes from the Online Mendelian Inheritance in Men (OMIM) database (Amberger et al., 2019). Overall, 1,113 diseases and 1,837 disease genes were associated to at least one affected tissue, with the majority associated to skeletal muscle (Fig. S1, Table S1).

**Figure 1.**
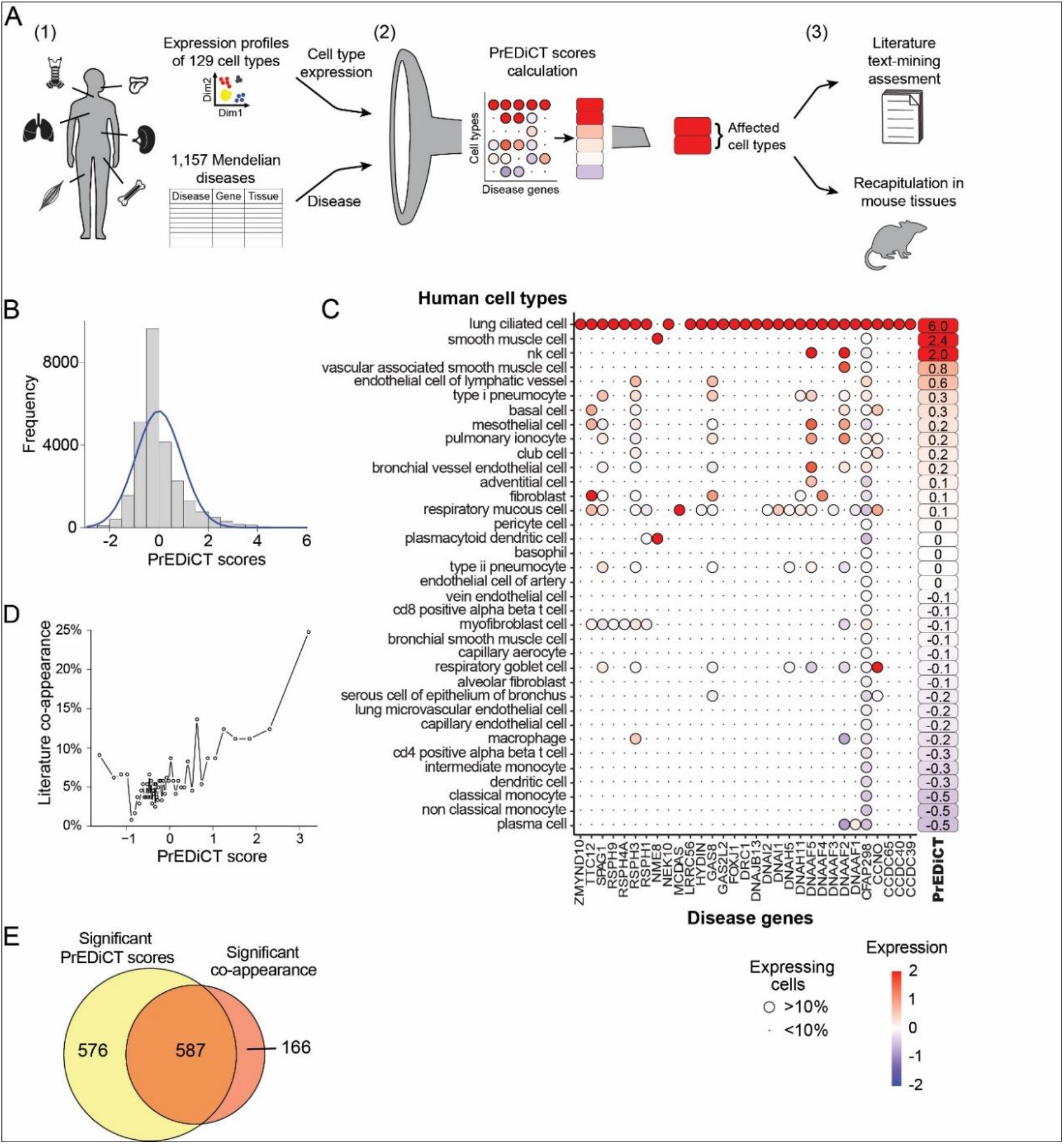
Overview of PrEDiCT calculation and assessment. A. The PrEDiCT workflow. In step 1, we analyzed single-cell expression data from six human tissues and 129 cell types, and associated 1,113 Mendelian diseases with their affected tissues. In step 2, we calculated the preferential expression of disease genes in cell types of disease-affected tissues, and used their median to produce the PrEDiCT score per disease and cell type. In step 3, we validated disease–cell-type associations with significant PrEDiCT scores via literature text-mining and analysis of mouse single-cell expression data. B. The distribution of PrEDiCT scores in human. C. The preferential expression of genes causal for primary ciliary dyskinesia (PCD) and the PrEDiCT scores of PCD in lung cell types. Preferential expression values and the percentage of cells expressing a gene are indicated by the color and the size of each circle, respectively. The resulting PrEDiCT score is displayed on the right and colored by the score value. D. The correlation between PrEDiCT scores of disease–cell-type pairs (X-axis) and the proportion of pairs with significant co-appearance in literature (Y-axis; r=0.5, p<0.001, Spearman correlation). Each dot represents 50 pairs. E. The overlap between the number of disease–cell-type pairs that obtained significant PrEDiCT scores (left circle) and the number of pairs with significant co-appearance in literature (right circle). The overlap was significant (p<E-15, Chi-squared test).

To calculate PrEDiCT scores, we computed the preferential expression of disease genes in each cell type relative to other cell types of the disease-affected tissue (Methods). Next, we set the PrEDiCT score of a disease in each cell type to the median preferential expression of the respective disease genes (Methods, Fig. S2). PrEDiCT scores ranged from −3 to 6, with a median of −0.3 (Fig. 1B). In particular, 1,163 disease–cell-type pairs had PrEDiCT scores ≥2, corresponding to the 95.6 percentile (top 5%), and were henceforth considered significant. An example of the ability of PrEDiCT to indicate affected cell types is presented by primary ciliary dyskinesia (PCD). PCD is characterized by damaged ciliary machinery in lung ciliated cells (Leigh et al., 2019) and was associated with 31 genes in our dataset. The highest preferential expression of most PCD-associated genes (29/31), and consequently the highest PrEDiCT score for PCD, was indeed obtained in lung ciliated cells (Fig. 1C).

To assess whether significant PrEDiCT scores indicate disease-affected cell types, we turned to the literature. We postulated that diseases and their affected cell types will co-appear in the literature more frequently than expected by chance. To estimate co-appearance, we searched PubMed for records that mentioned a disease or a cell type of disease-affected tissues by using Biopython package [(Cock et al., 2009); Table S2]. Next, we tested the significance of the co-appearance of all possible disease–cell-type pairs (Methods). Overall, 753 pairs co-appeared in the literature significantly more often than expected by chance (adjusted p<0.001, Chi-squared test). PCD, for example, co-appeared significantly with lung ciliated cells, as well as with respiratory goblet, mucous, and basal cells (the latter differentiating to the other three). Notably, PrEDiCT scores of all disease–cell-type pairs and the probability of their co-appearance were correlated (r=0.5, p<0.001, Spearman correlation; Fig. 1D). In particular, the 1,163 pairs with significant PrEDiCT scores were enriched for pairs with significant co-appearance in the literature (18%, expected 6%, p<E-15, Chi-squared test; Fig. 1E), and were henceforth considered as affected. In total, 714/1,113 diseases were determined as affecting 110/129 cell types, creating a uniquely large resource of disease-affected cell types (Table S3).

### Disease-affected cell types are recapitulated in mouse

To further assess whether PrEDiCT scores indicate disease-affected cell types, we tested whether similar cell types were affected in mice. For this, we downloaded mouse single-cell transcriptomes for the six tissues from Tabula Muris (Tabula Muris, 2018). These data consisted of 46 annotated cell types and some unannotated subsets (Fig. 2A).

**Figure 2.**
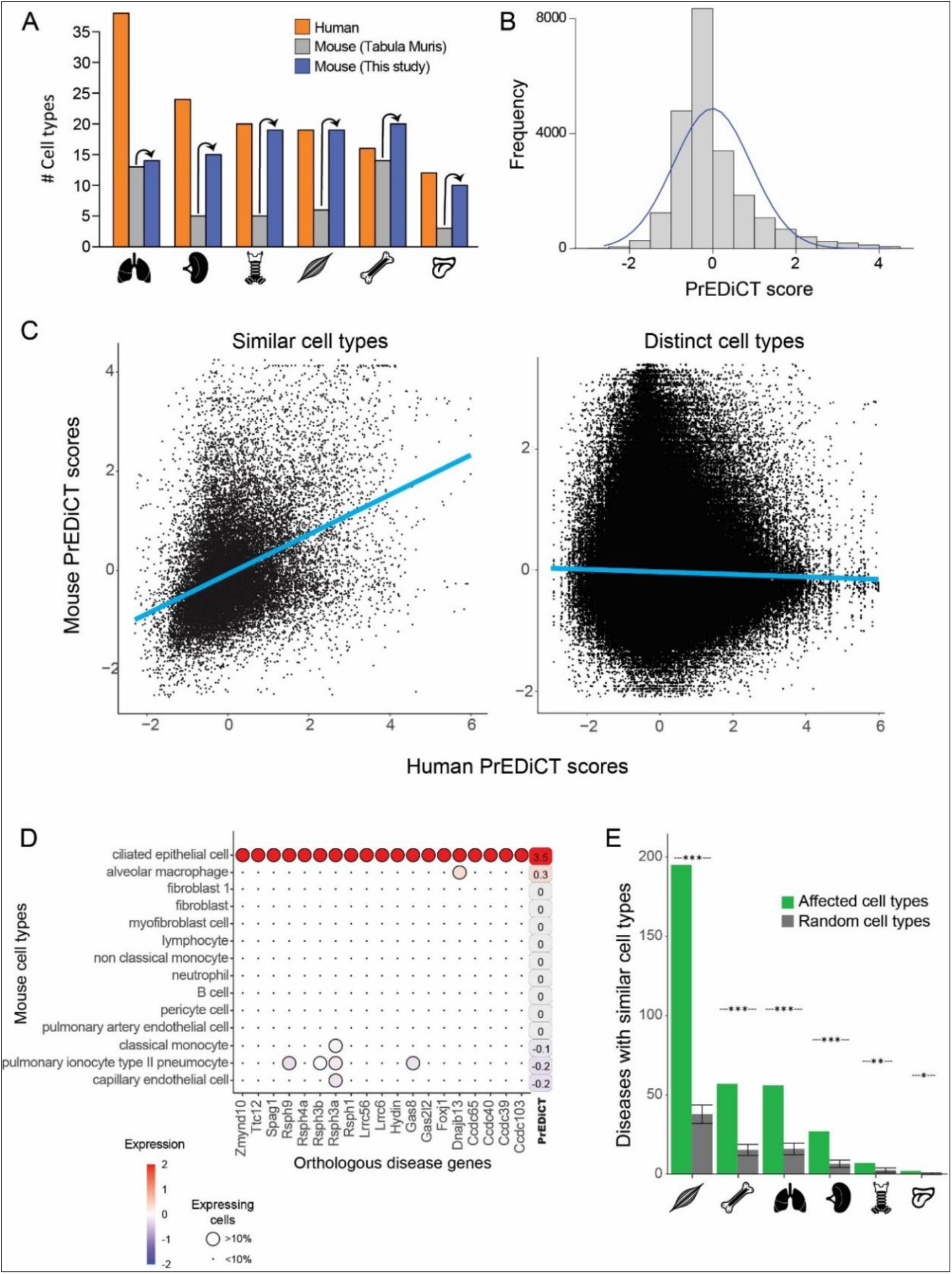
Recapitulation of disease-affected cell types in mouse. A. The number of human cell types annotated by Tabula Sapiens [(Tabula Sapiens, 2022); red], and the number of mouse cell types annotated by Tabula Muris [(Tabula Muris, 2018); grey] and this study (blue). B. The distribution of PrEDiCT scores in mouse. C. The correlation between PrEDiCT scores in human (X-axis) and mouse (Y-axis) cell types. Each dot represents a distinct pair. PrEDiCT scores of distinct cell types did not correlate (left; r=-0.02, Spearman correlation), in contrast to PrEDiCT scores of similar cell types (right; r=0.39, p<E-15). D. The preferential expression of mouse orthologs of PCD disease genes and the PrEDiCT scores of PCD in mouse lung cell types. Preferential expression values and the percentage of cells expressing a gene are indicated by the color and the size of each circle, respectively. The resulting PrEDiCT score is indicated on the right and colored by the score value. E. The cell types affected by the same disease in human and mouse were more likely to be similar (green) than expected by chance (grey). Adjusted *p<0.05, **p<0.01, ***p<0.001, permutation test.

To improve the annotation of cell types, we reanalyzed the single-cell mouse transcriptomes (Methods). Altogether, we obtained 97 cell clusters across all six tissues (Table S4). Sixteen clusters overlapped considerably with cell types annotated by Tabula Muris and were thus annotated similarly. We annotated the 81 remaining clusters by careful examination of the expression of known cell-type marker genes (Methods), thereby improving cell type annotation per tissue (Fig. 2A). For instance, the number of annotated cell types in skeletal muscle increased from six to 19. Newly annotated cell types included clinically relevant cell types, such as basement-membrane residing fibroblasts that are a main source of different collagens and are essential for skeletal muscle physiology (Kivirikko et al., 1995; Zou et al., 2008). Next, we matched between similar human and mouse cell types based on expression of orthologous marker genes, by using matchSCore2 package [Methods; (Mereu et al., 2020)]. As expected, a large variety of human cell types were similar to mouse cell types (82/129) and vice versa (70/97, Table S5).

Lastly, we tested whether diseases affected similar cell types between human and mouse (Fig. 2A, Fig. S3). For this, we calculated PrEDiCT scores in mouse cell types, based on expression of mouse orthologs of human disease genes. The distribution of PrEDiCT scores was similar between mouse and human (Fig. 2B; Table S6). As with human data, PrEDiCT scores ≥2 ranked at the 95.5 percentile (top 5%), resulting in 1,036 significant pairs of diseases and mouse cell types. We compared the PrEDiCT scores of all cell-type pairs of the corresponding tissues between the species. Whereas PrEDiCT scores of distinct cell types did not correlate (r=-0.02, Spearman correlation), PrEDiCT scores of similar cell types were modestly correlated (r=0.39, p<E-15, Spearman correlation; Fig. 2C). Compatible with our previous proof-of-concept, PCD scored highest in mouse ciliated epithelial cells, which were determined to be similar to human lung ciliated cells (Fig. 2D). Of the 714 diseases with affected cell types in humans, 303 diseases (42%) affected similar cell types in mice (Table S1), a fraction that was larger than expected by chance (Fig. 2E; p<E-4, permutation test; Methods). This enrichment, and the observation that commonly affected cell types were not biased toward specific cell types, support the validity and generality of the PrEDiCT scheme.

### Characteristics of disease-affected cell types

Are certain cell types more likely to be affected by hereditary disease? To answer this question, we first scored the susceptibility of each cell type. Susceptibility score of a given cell type was set to the fraction of diseases affecting that cell type out of all diseases that affect its tissue. Susceptibility scores ranged from 0% to 21% (Table S7). Most susceptible were Schwann cells of the tongue (scored 21%), which were affected mostly by neurogenic diseases, such as ‘X-linked infantile spinal muscular atrophy 2’ that manifests with tongue fasciculations.

Then, we asked whether the prevalence of a cell type in a tissue correlated with its susceptibility. For that, we compared between cell-types prevalence in a tissue, as determined by the fraction of corresponding cells within the tissue, and susceptibility scores (Table S7). The two measures did not correlate (r=-0.09, Spearman correlation; Fig. 3A). For example, ionocytes were the most susceptible cell type in lung and the second highest in trachea, while accounting for less than 1% of the cells in those tissues.

**Figure 3.**
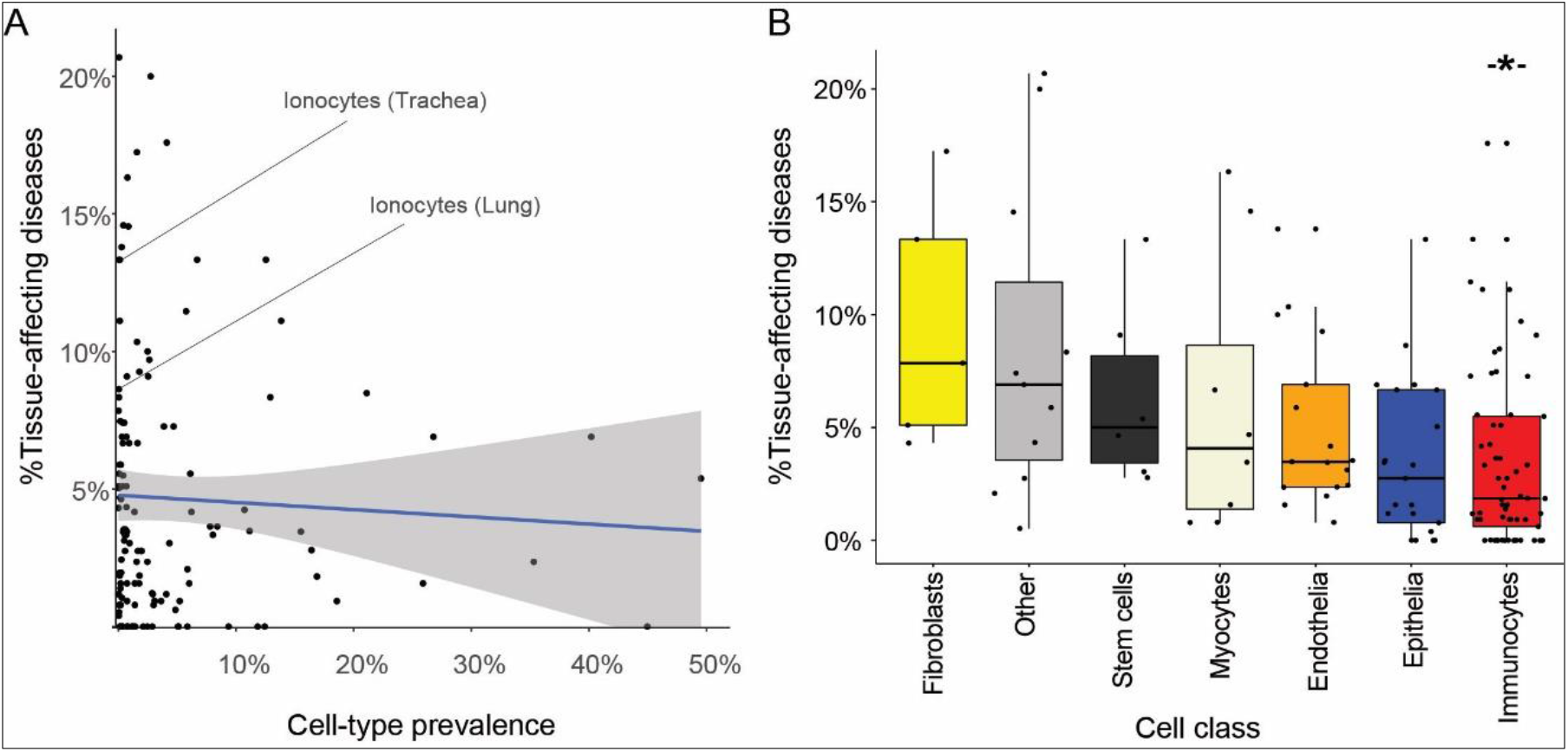
Characteristics of disease-affected cell types. A. The tendency of cell types to be affected by hereditary diseases (Y-axis) was not correlated with their tissue prevalence (X-axis). The blue line represents linear correlation (r=-0.09, p=0.33, Spearman correlation). B. The tendency of cell-type classes to be affected by hereditary diseases varied between classes (p=0.002, ANOVA test). Immunocytes were least susceptible (adjusted p=0.019, Mann-Whitney U test, Methods).

Lastly, we assessed the susceptibility of cell classes. For that, we classified cell types as fibroblasts, stem cells, myocytes, endothelia, epithelia, immunocytes and other. Next, we associated each cell class with the median susceptibility scores of its cell types (Table S7). Cell classes had varying susceptibility scores (Fig. 3B; p=0.002, ANOVA), with immunocytes being least susceptible (adjusted p=0.019, Mann-Whitney U test, Methods). Fibroblasts, in contrast, were the most susceptible, yet with borderline significance (adjusted p=0.056), potentially due to their low number of cell types.

### Diseases with multiple inflicted tissues affect similar cell types

Most diseases in our dataset manifested clinically in a single tissue, in accordance with previous observations (Hekselman & Yeger-Lotem, 2020). Yet, 30% (217/714) of the diseases manifested in two or more tissues, and were denoted multi-tissue diseases (Fig. 4A). We suspected that these diseases affect a cell type that is similar between the disease-manifesting tissues. To test this hypothesis, we identified similar cell types between tissues, and then examined whether cell types affected by the same disease were enriched for similar cell types. Similar cell types were identified using matchSCore2 package [Methods; (Mereu et al., 2020)]. Overall, we identified 840 pairs of similar cell types between tissues, of which 52% were immunocytes, including macrophages, neutrophils, T cells, B cells, as well as endothelia and fibroblasts. Next, we examined whether the cell types affected by the same disease were enriched for similar cell types. We found that 36% (78/217) of the diseases affected at least one pair of similar cell types, a fraction that was about 4-fold higher than expected by chance (Fig. 4B, Fig. S4; adjusted p<0.05, permutation test; Methods). For instance, chronic granulomatous disease, which results in splenomegaly and pneumonia due to impaired neutrophils (Leiding & Holland, 1993), affected neutrophils in both spleen and lung (Fig. 4C).

**Figure 4.**
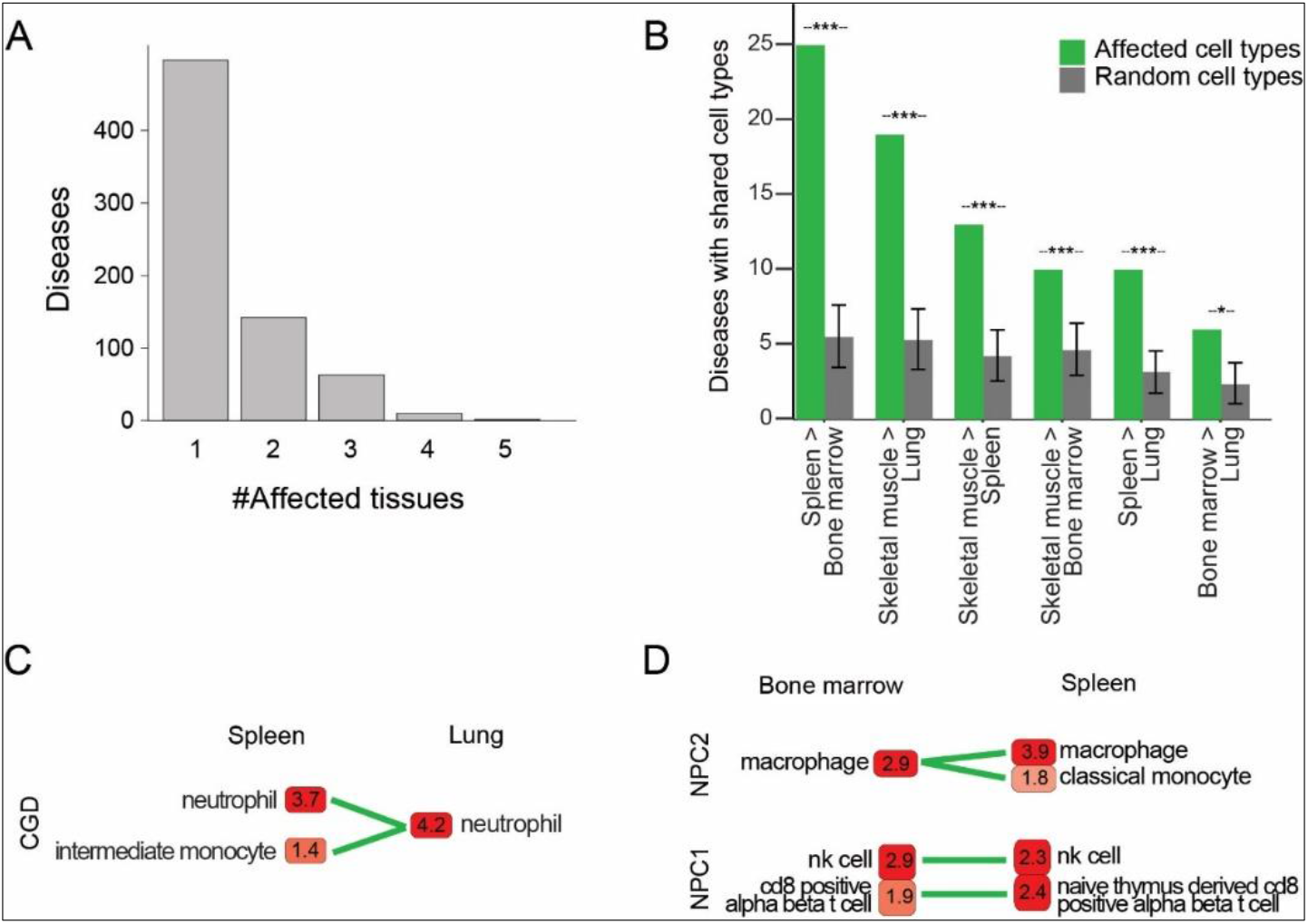
Multi-tissue diseases tend to affect similar cell types in those tissues. A. The numbers of diseases (Y-axis) grouped by the number of affected tissues (X-axis). B. For each disease, affected cell types in one tissue were shared with another affected tissue (green), more than expected by chance (grey; Permutation test; Methods). Under the bars, tissue (first) and the one it is compared to (second) are shown. Only non-redundant tissue-pairwise comparisons sharing >5 affecting diseases are shown. All tissue-pairwise comparisons are shown in Fig. S4. Adjusted *p<0.05, ***p<0.001. C. PrEDiCT scores of cell types affected by chronic granulomatous disease (CGD) in spleen and lung. Shared cell types were connected by green lines. D. PrEDiCT scores of cell types affected by Niemann-Pick disease type C1 (NPC1; bottom) and type C2 (NPC2; top) in bone marrow and spleen. Shared cell types were connected by green lines.

Intriguingly, some multi-tissue diseases did not affect similar cell types. For example, Niemann-Pick disease type C (NPC) is a lipid storage disorder characterized mainly by progressive neurodegeneration. The two types of NPC, NPC1 and NPC2, which are caused by aberration in distinct genes, affected distinct cell types (Vance, 2006). NPC2 affected macrophages residing in bone marrow and spleen, in agreement with what was previously known for NPC (Bajwa & Azhar, 2022). In contrast, NPC1 affected cytotoxic lymphocytes in both tissues (specifically natural-killer and CD8 T cells; Fig. 4D). This finding, which has yet to be established experimentally, is compatible with the functional impairment of the cytotoxic lymphocytes seen specifically in NPC1 patients (Castiblanco et al., 2022; Speak et al., 2014).

### Disease genes are a central component in the expression signature of cell types

As preferential expression of disease genes marks affected cell types, could their expression pattern distinguish one cell type from another? To test this, we clustered cell types by the expression of disease genes and by the expression of all genes. Cell types of the same cell-type class tended to cluster together according to both datasets. Next, we compared between the resulting cell-type hierarchy (dendrogram) per tissue, by using dendextend R package [(Galili, 2015); Methods]. In all tissues, the two dendrograms were similar to each other (median Cophenetic correlation=0.8, Table S8) more than expected by chance (adjusted p<0.05, permutation test). Similarity was prominent even in skeletal muscle and lung tissues, which had the widest variety of cell-type classes (Cophenetic correlation=0.9 and 0.7, respectively, adjusted p<0.05; Fig. 5). Altogether, these results demonstrate that the expression signature of disease genes is a central component in the identity of cell types and cell classes.

**Figure 5.**
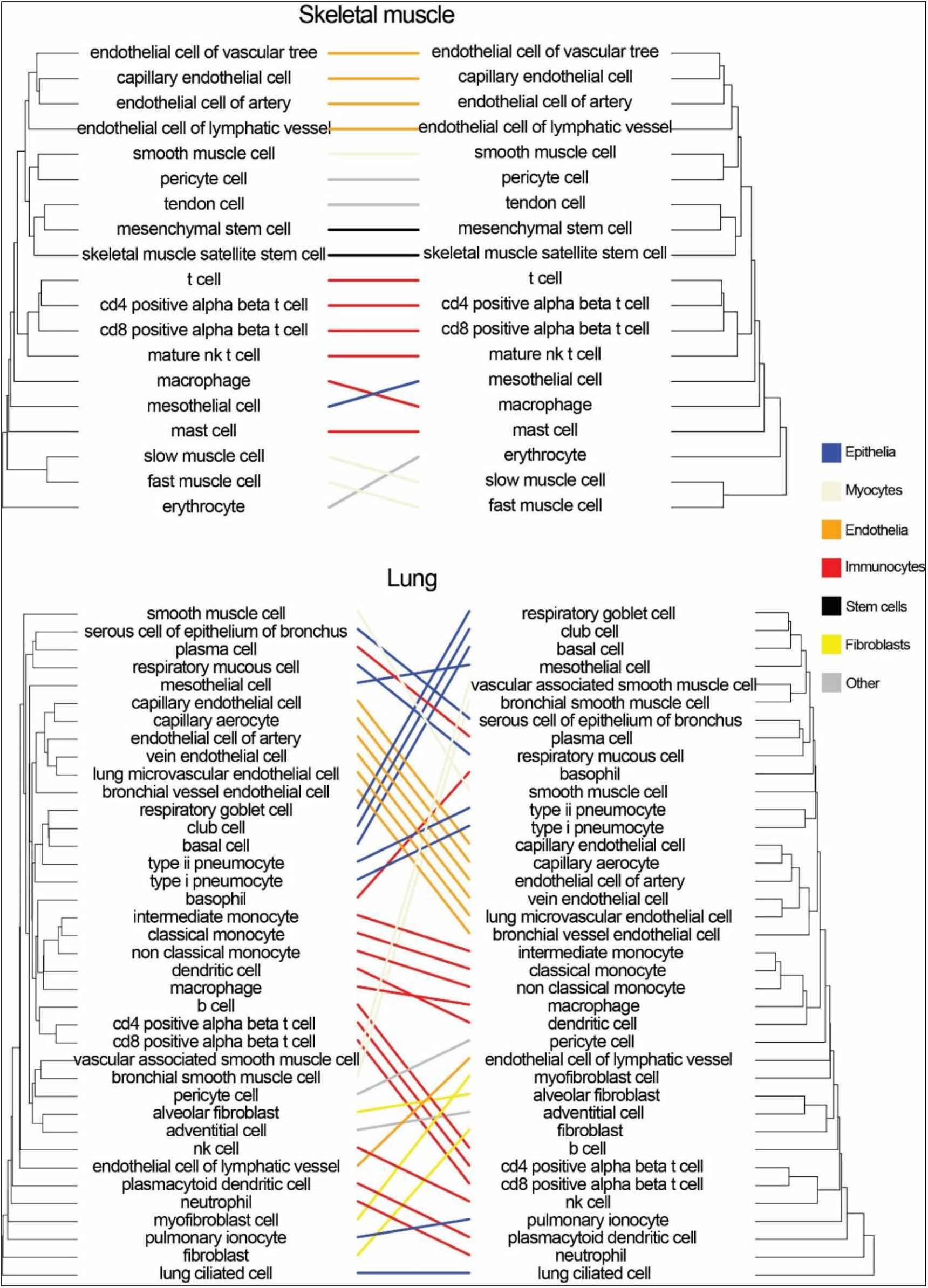
Cell-type dendrograms based on disease genes (left) or all genes (right) in skeletal muscle and lung tissues. Lines connect the corresponding cell types between dendrograms and are colored by cell class. Cophenetic correlation was 0.9 and 0.7, respectively, adjusted p<0.05, permutation test.

### Cell types affected by mitochondrial diseases and heritable cancers

Heritable mitochondrial diseases may manifest clinically in various tissues, depending on the type of defect and stress incurred (Pacheu-Grau et al., 2018). To examine the cell types affected by heritable mitochondrial diseases, we manually extracted 15 mitochondrial diseases and their 62 causal genes from OMIM (Table S9). The 15 mitochondrial diseases manifested in skeletal muscle, as well as bone marrow (four diseases), lung (two diseases), and spleen (two diseases). Next, we calculated PrEDiCT scores per cell type in those tissues. In skeletal muscle, we identified affected cell types for 12/15 mitochondrial diseases (Fig. 6A). Compatible with the high demand for mitochondrial activity in muscle cells, ten of these diseases affected both slow and fast muscle cells, e.g., mitochondrial complex I, II, III, IV and V deficiencies (OMIM: PS252010, PS252011, PS124000, PS220110 and PS604273, respectively). Additionally, progressive external ophthalmoplegia with mtDNA deletions (OMIM: PS157640) was scored highest in slow and fast skeletal muscles, yet with borderline significance (score 1.9 and 1.8). In lung, mitochondrial complex I and V deficiency (OMIM: 619003 and 615228) affected ionocytes. Intriguingly, both diseases are characterized by lung hypoplasia, suggesting a role of ionocytes in lung development.

**Figure 6.**
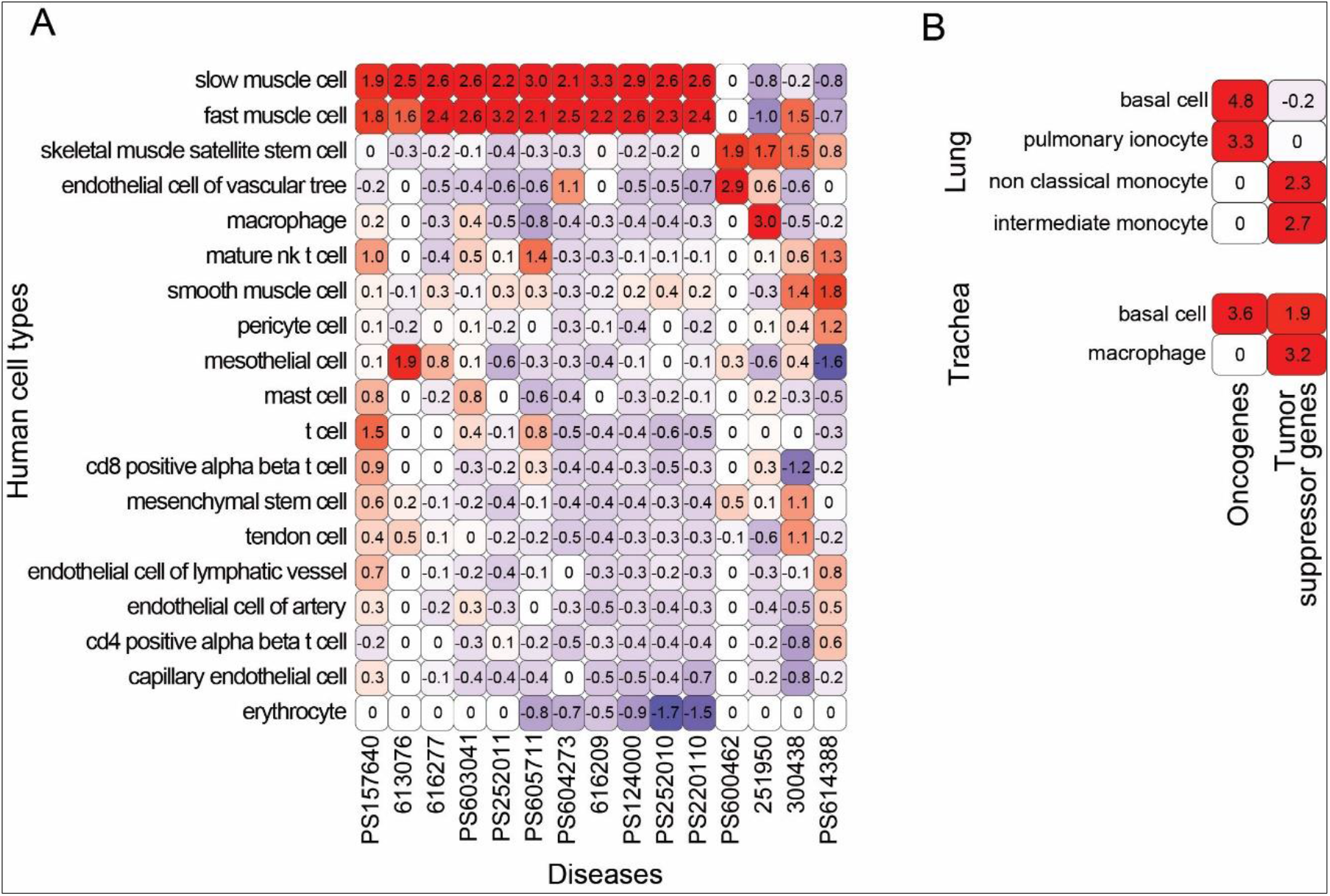
Cell types affected by mitochondrial diseases and heritable cancers. A. PrEDiCT scores of mitochondrial diseases in skeletal muscle cell types. Diseases and cell types were clustered according to their Euclidean distance. PS157640 = progressive external ophthalmoplegia with mtDNA deletions; 613076 = myopathy, mitochondrial progressive, with congenital cataract, hearing loss, and developmental delay; 616277 = mitochondrial short-chain enoyl-coa hydratase 1 deficiency; PS603041 = mitochondrial DNA depletion syndrome; PS252011 = mitochondrial complex II deficiency, nuclear type; PS605711 = multiple mitochondrial dysfunctions syndrome; PS604273 = mitochondrial complex V (ATP synthase) deficiency, nuclear type; 616209 = myopathy, isolated mitochondrial, autosomal dominant; PS124000 = mitochondrial complex III deficiency, nuclear type; PS252010 = mitochondrial complex I deficiency, nuclear type; PS220110 = mitochondrial complex IV deficiency, nuclear-type; PS600462 = myopathy, lactic acidosis, and sideroblastic anemia; 251950 = mitochondrial myopathy with lactic acidosis; 300438 = Hsd10 mitochondrial disease; PS614388 = encephalopathy due to defective mitochondrial and peroxisomal fission. B. PrEDiCT scores of cell types affected by heritable cancers in lung (top) and trachea (bottom), based on the preferential expression of oncogenes (left) and tumor suppressor genes (right).

Heritable cancers often show tissue-selective manifestation (Bianchi et al., 2020). To examine cancer-affected cell types, we retrieved 25 heritable cancers, their oncogenes and their tumor suppressor genes from the Cancer Gene Census [Methods; (Sondka et al., 2018)]. We manually annotated cancer-manifesting tissues, and then calculated PrEDiCT scores per cell type according to the expression of either oncogenes or tumor suppressor genes (Table S10). In both lung and trachea, PrEDiCT scores based on oncogenes were significant in basal cells (Fig. 6B). PrEDiCT scores based on tumor suppressor genes were significant in the monocytic lineage in both tissues.

### Disease-related intercellular communication routes

Important intercellular communication routes could be revealed by focusing on diseases caused by aberrant ligands or receptors. We retrieved diseases that were associated with at least two genes encoding a ligand and its receptor, and were upregulated in distinct cell types (Methods). We collected six such diseases involving 12 ligand-receptor pairs. For instance, autosomal recessive limb-girdle muscular dystrophy has four ligand-encoding disease genes (collagens COL6A1, COL6A2, COL6A3 and laminin LAMA2) and a single receptor-encoding disease gene (alpha-dystroglycan, DAG1). All ligands were upregulated in mesenchymal stem cells, whereas the receptor was upregulated in satellite stem cells. This suggests that the disease disrupts communication signals from mesenchymal stem cells to satellite stem cells (Fig. 7A). In support of this model, disruption of DAG1 in satellite stem cells has been associated with the defective muscle regeneration seen in these patients (Cohn et al., 2002; Servian-Morilla et al., 2020). Additionally, the disease was attenuated in ligand-knockout [Col6a1(-/-)] model mice by supplying wild-type mesenchymal-stem-cell derived fibroblasts. This treatment not only restored the collagen VI of the tissue, but also rescued defects in satellite stem cells (Urciuolo et al., 2013).

**Figure 7.**
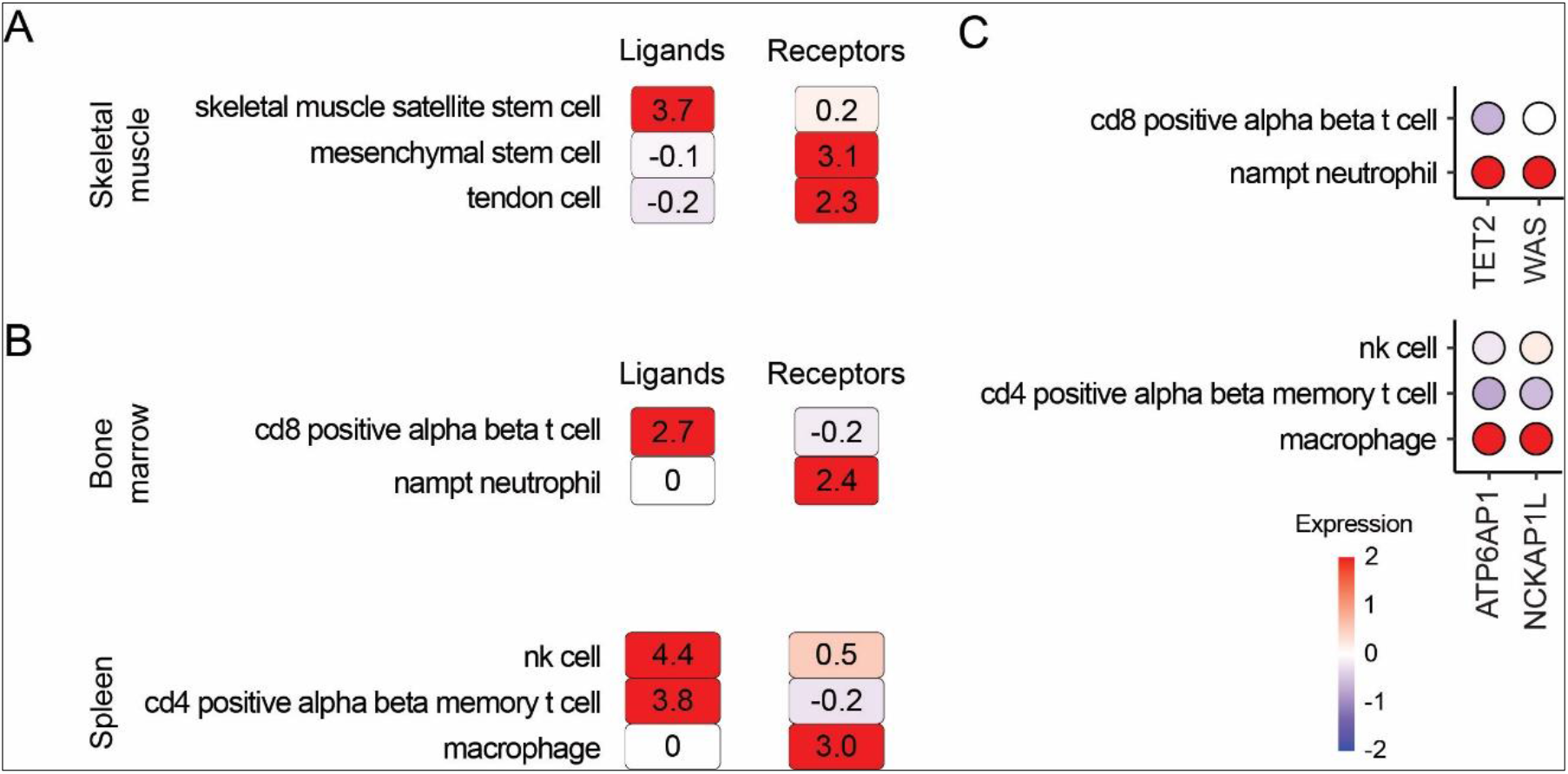
Diseases that disrupt intercellular communication affect distinct cell types. A. The PrEDiCT scores of autosomal recessive limb-girdle muscular dystrophy in skeletal muscle. PrEDiCT scores were based on the expression of disease genes encoding ligands (left) and receptors (right). Only cell types with significant PrEDiCT scores are shown. B. The PrEDiCT scores of immunodeficiency disease in bone marrow (top) and spleen (bottom), as described in A. C. The expression of disease genes causal for immunodeficiency, which were upregulated in the cell types affected by receptor aberrations. In the balloon plot, the gene expression values are indicated by the color of each circle, as indicated in the key. Genes were expressed in ≥10% cells of each cell type.

Another interesting case is immunodeficiency disease that has three ligand-encoding disease genes (LCK, IFNG and CD40LG) and their receptor-encoding disease gene (interferon-gamma receptor 1, IFNGR1). The three ligands were upregulated in T cells, whereas IFNGR1 was upregulated in myeloid phagocytes (neutrophils and macrophages in bone marrow and spleen, respectively; Fig. 7B). This was compatible with the well-studied immune-related responsiveness of myeloid phagocytes via IFNGR1 (Cabral-Marques et al., 2017; Ellis & Beaman, 2004; Eshleman et al., 2020). Also, Aberration in disease genes other than ligands and receptors might also disrupt intercellular communication routes by acting downstream of the receptor. Indeed, by examining the causal genes for immunodeficiency that were upregulated in myeloid phagocytes, we found TET2 and WAS upregulated in bone marrow, and ATP6AP1 and NCKAP1L upregulated in spleen. Intriguingly, all four disease genes are known to participate in signaling pathways associated with myeloid-phagocytes–mediated immunity (Banks et al., 2021; Brunetti et al., 2022; Stahnke et al., 2021; Suwankitwat et al., 2021; Yang et al., 2020). These findings suggest that disease genes that are upregulated in the receptor-bearing cell type might be involved in a downstream signaling pathway.

## DISCUSSION

Here, we presented the PrEDiCT method for identifying disease-affected cell types based on the preferential expression of Mendelian disease genes. Preferential expression of disease genes was shown to characterize disease-affected tissues in many diseases (Barshir et al., 2018; Barshir et al., 2014; Hekselman & Yeger-Lotem, 2020; Lage et al., 2008; Sonawane et al., 2017). Also, it has been applied recently to highlight potentially-affected cell types for several diseases (Dai et al., 2021; Eraslan et al., 2022; Jagadeesh et al., 2021; Kim-Hellmuth et al., 2020; Zhao et al., 2021). However, validation was rarely conducted (Montoro et al., 2018; Plasschaert et al., 2018). Here, in contrast, we corroborated the relevance of PrEDiCT by showing that significantly high PrEDiCT scores were more likely supported by literature (Fig. 1) and recapitulated in mouse (Fig. 2).

The use of preferential expression to highlight affected cell types has limitations. Apart from being oblivious to post-translational regulation, most available single-cell transcriptomic datasets do not contain full-length gene reads and thus cannot be used to assess alternatively spliced transcripts. Additionally, preferential expression is a noisy measure, which might lead to erroneous associations of diseases to their affected cell types. For example, disease genes were shown to be upregulated in cell types that reside in unaffected tissues (Eraslan et al., 2022; Jagadeesh et al., 2021). For instance, both muscle and breast adipocytes upregulated metabolic-myopathy–associated genes, yet only the muscle seems to be affected by the disease (Eraslan et al., 2022). To reduce irrelevant associations, we studied diseases only in the context of their affected tissues (Fig. S1). To enhance relevant associations, the PrEDiCT of diseases with multiple disease genes were based on the combined scores of these genes. As we showed for PCD, the combined score indeed pointed to PCD-affected cell types (Fig. 1C). Lastly, preferential expression is just one of several mechanisms that lead to tissue-selective disease manifestations (Hekselman & Yeger-Lotem, 2020). Nevertheless, by applying PrEDiCT to 1,113 diseases and single cell transcriptomes of six distinct tissues, we revealed affected cell types for 64% of the Mendelian diseases in our dataset (Table S3). This fraction is similar to the fraction observed for Mendelian diseases in tissue contexts (Barshir et al., 2018; Barshir et al., 2014; Hekselman & Yeger-Lotem, 2020; Lage et al., 2008; Sonawane et al., 2017).

Altogether, 714 diseases affected 110 out of the potential 129 cell types in those tissues. Interestingly, there was no correlation between cell type prevalence and its likelihood to be affected (Fig. 3A). In particular, immunocytes were the least likely to be affected (Fig. 3B). This suggests that immunocytes are either more resilient than other cell types, or, alternatively, that their impairment is lethal to the organism. Notably, a cross-tissue study of 20 human tissues showed that immunocytes have similar expression signatures across tissues, in accordance with their common functions, whereas endothelia have tissue-specific expression signatures that reflect their tissue-specialized roles (Tabula Sapiens, 2022). We found that endothelia were affected by diseases like other cell classes. Hence, it seems that germline impairment of immunocytes is more likely lethal, whereas the tissue-specialization of endothelia limits the impact of their germline impairment and thus facilitates overall survival.

Our expansive resource of diseases and affected cell types can be used to interrogate disease etiologies. For example, we found that mitochondrial diseases tend to affect muscle cells, in accordance with their energetic demands (Fig. 6A). Likewise, in familial cancers, oncogenes and tumor suppressor genes were associated with distinct cell types, specifically with basal cells and with monocytes, respectively (Fig. 6B). This suggests that protective cells, such as monocytes, are more susceptible to mutation in tumor suppressor genes, whereas tissue-constructive cells are more susceptible to mutations in oncogenes. This corresponds with the function of monocytes in the elimination of malignant transformation of cells in different tissues (Robinson et al., 2021). Another interesting subset of diseases are those that impair intercellular communication. A recent study explored the cell types affected by monogenic muscular disorders (Eraslan et al., 2022). In agreement with their results, we found that autosomal recessive limb-girdle muscular dystrophy disrupts intercellular communication among muscle cell types, via mutations in the DAG1 receptor (Fig. 7). However, by interrogating the preferential expression of four disease-associated ligands of DAG1, we also highlighted the involvement of mesenchymal stem cells in the disease (Urciuolo et al., 2013). Additionally, disease genes that are upregulated in the receptor-bearing cell types could be part of a downstream disease-associated pathway. For instance, owing to upregulation of IFNGR1 gene, which encodes an immunodeficiency-associated receptor, we found myeloid phagocytes as affected by this disease. Then, we found three additional disease genes that were upregulated in myeloid phagocytes and were all associated with myeloid-phagocytes–mediated immune response (Banks et al., 2021; Cabral-Marques et al., 2017; Ellis & Beaman, 2004; Eshleman et al., 2020; Stahnke et al., 2021; Suwankitwat et al., 2021; Yang et al., 2020). Thus, exploring disease genes in appropriate cellular context can enhance the mechanistic understanding of disease emergence.

The associations between diseases and affected cell types, though supported by literature and recapitulated in mice, remain putative. Experimental testing could be performed in human cell lines, or in mouse models, in light of the many shared cell types between human and mouse (Fig. 2). These validation experiments have a huge potential to open new directions in disease research and accelerate cell-directed gene therapy.

## METHODS

### Human single-cell transcriptomics analysis

Single-cell transcriptomes were downloaded from Tabula Sapiens (Tabula Sapiens, 2022). We focused on tissues that consisted of ≥2 samples with ≥800 cells and were sequenced in both human and mouse using microfluidic droplet-based 3’-end technology. These tissues included bone marrow, lung, skeletal muscle, spleen, tongue, and trachea. Analysis was done using Seurat package v4.0.5 (Hao et al., 2021). Per tissue, gene expression levels were normalized cell-wise using the NormalizeData function. Henceforth, we considered only genes with normalized counts ≥0.05 in ≥10% of cells of at least one cell type. Their average expression was calculated using AverageExpression function.

### Mouse single-cell transcriptomics analysis

Single-cell transcriptomes were downloaded from Tabula Muris (Tabula Muris, 2018). To improve the annotation of mouse cells from Tabula Muris, we reanalyzed the transcriptomic profiles of each tissue. First, we selected 2,000 variably expressed genes using FindVariableFeatures function in Seurat. The minimum and maximum average normalized expression of genes across cells were set to 0.05 and 3, respectively (mean.cutoff=c(0.05,3)). We scaled and centered the expression values of the variably expressed genes using ScaleData function, while correcting for the different samples (vars.to.regress=‘mouse.id’). Then, we projected their expression on all significant principal components (PCs; p<0.001, JackStraw test) ordered by their explained variance.

Next, we applied a two-phase clustering process. We clustered cells using Seurat FindNeighbors and FindClusters functions based on all the top significant PCs. To resolve over-clustering of cells, we hierarchically ordered cell clusters using BuildClusterTree function. Then, we tested whether cells from different splits in the tree were distinguishable, according to out-of-bag error of a random forest classifier that was trained on variably expressed genes. Indistinguishable cell clusters (p≥0.05) were merged. To estimate sample-based differences between related clusters, we repeated hierarchically ordering of merged cell clusters. Terminal splits of cell clusters that included uneven numbers of cells from different samples (adjusted p<0.001, Chi-square) were merged. Such differences were observed only in tongue and lung tissues.

### Annotations of mouse cell clusters

We compared the clusters obtained above to the cell type annotations of Tabula Muris. A cluster where a similar annotation was common to >90% of its cells, and where >90% of cells with that annotation were within that cluster, was annotated according to Tabula Muris. All other clusters were manually annotated based on highly expressed marker genes (Z-score ≥ 2). We manually searched PubMed for evidence that any of these markers indicates a known cell type (including cell identity and\or function), preferably in the context of the relevant tissue. A cluster was annotated if at least two of its markers indicated the same cell type. To comply with other studies, cell types were named as in Cell Ontology (Diehl et al., 2016). Cell type annotations, relevant marker genes and supporting literature appear in Table S4.

### Diseases-tissue annotations and curation

Disease data were retrieved from OMIM (Amberger et al., 2019), and included diseases (i.e., phenotypes with a known molecular basis and phenotypic series). Each disease was associated with its disease genes and their mouse orthologs according to OMIM, and was associated with its affected tissue according to HPO (Kohler et al., 2021). We focused on disease that were cataloged by HPO as having main phenotypic abnormality in blood and blood-forming tissues (HP: 0001871), lungs (HP: 0002088), musculature (HP: 0003011), spleen (HP: 0001743), tongue (HP: 0000157) and trachea (HP: 0002778), in accordance with the six tissues that we analyzed. Phenotypic abnormalities that HPO categorized under each of these main terms were also included. We assessed the compatibility of associations by comparing them to ODiseA database, which includes manually curated associations of diseases and their affected tissues (Hekselman et al., 2022). For this, we downloaded from ODiseA all the diseases that affected blood and bone marrow, lung, and trachea. Diseases that were not supported by ODiseA were excluded from further analysis.

### PrEDiCT score calculation

Per tissue, we calculated the PrEDiCT scores based on the average expression of genes across cell types. We included only genes expressed above the median average expression in that tissue. We applied a Z-score transformation to the average expression of each gene across cell types. The preferential expression of a gene in a cell type was set to its Z-score in that cell type. Then, the PrEDiCT score of a disease in a cell type was set to the median preferential expression of its disease genes.

### Text-mining of PubMed records

We searched PubMed records for publications containing names of the disease in our disease dataset or names of human cell types in the disease-affected tissues. For this, we used eSearch function in Biopython package (Cock et al., 2009), with number of maximum publications retrieved set to 100,000 (retmax=100,000) and all other parameters set to default. Then, per tissue, we intersected the list of publications of each tissue-affecting disease *d* with the list of publications of each tissue cell type *c*, to identify publications that mention both. Diseases with less than three publications that mentioned it together with a specific cell type were excluded. Cell types that were not mentioned with any disease were excluded. Next, per tissue, we determined whether disease–cell-type pairs were co-appearance significantly by applying a Chi-squared test. Pairs with Z scores higher than expected (adjusted p<0.001, Bonferroni correction), were determined as significant.

### Determining similar cell types between tissues

For each pair of tissues, all cell types were compared to each other. For this, each cell type was associated with marker genes, namely genes with Z-score ≥2. Markers of mouse cell types were converted to their human orthologous genes according to the Mouse Genome Informatics database (Bult et al., 2019). Next, we estimated the similarity between each pair of cell types using matchSCore2 package, which compared the two markers lists by using Jaccard index (Mereu et al., 2020). Cell types were determined as similar if their Jaccard index was ≥0.05 and in the top 10th percentile.

### Permutation tests for similarity between affected cell types

For each pair of tissues, we denoted one as the reference tissue (*Tr*) and the other as the test tissue (*Tt*). Success was determined, per disease, if any of the cell types in *Tt* were similar to any of the affected (PrEDiCT score ≥ 2) cell types in *Tr*. We tested the null hypothesis that the number of diseases for which success was determined (*num_s*) was not higher than expected by chance. For this, we carried out a permutation test. Per disease, we randomly selected cell types equal in number to the affected cell types in *Tr* and repeated the success test; specifically, we checked if the randomly selected cell types in *Tr* were similar to any of the affected cell types in *Tt*. We repeated this for all diseases with affected cell types in *Tr*, and recorded the number of randomly successful diseases (*num_r*). We repeated this procedure 1,000 times. Significance was calculated as the fraction of cases where num_r ≥ num_s. P-values were adjusted for multiple comparisons by Benjamini-Hochberg correction.

### Analysis of affected cell classes

Susceptibility score of a given cell type was set to the fraction of diseases affecting that cell type out of all diseases that affect that same tissue (Table S7). Cell types were grouped into one of five cell classes: fibroblasts, stem cells, myocytes, endothelia, epithelia and immunocytes, or were grouped as ‘other’. Then, we applied an analysis of variance (ANOVA) test to the proportions of diseases that affected each cell class using aov function in R v4.1.1. To further determine whether specific cell classes were more susceptible than others, we compared the susceptibility scores of cell types from each class to all other cell types, by using Mann-Whitney U test. P-values were adjusted for multiple comparisons by Benjamini-Hochberg correction.

### Comparison between cell-type dendrograms

Dendrograms of cell types in each tissue were compared using dendextend v1.15.2 in R (Galili, 2015). First, as recommended by dendextend, distance matrices were computed based on the Euclidean distance of the average expression profiles of cell types. Next, cell types were hierarchically clustered using unweighted pair group method with arithmetic mean (UPGMA). The entire process was applied twice: (a) based on disease genes, and (b) based on all expressed genes. Using untangle function, both dendrograms were visualized. Similarity between two dendrograms, denoted cophenetic distance, was calculated using cor_cophenetic function. Lastly, significance of cophenetic distance was calculated by permuting over the labels of the second dendrogram (i.e., based on all expressed genes) 1,000 times, and assessing the distribution under the null hypothesis. P-values were adjusted for multiple comparisons by Benjamini-Hochberg correction.

### Heritable cancers

Data of heritable cancers were downloaded from the Cancer Gene Census of the Catalogue of Somatic Mutations in Cancer [COSMIC; (Sondka et al., 2018)]. Specifically, we downloaded germline tumor type, associated genes, and role in cancer (oncogenes or tumor suppressor genes) from tiers 1 and 2. We manually annotated germline tumor types to the six tissues included in our study.

### Ligands- and receptors-associated diseases

We extracted a list of 1,625 pairs of ligands and their corresponding receptors from (Jin et al., 2021). We retrieved all diseases with disease genes that included both a ligand and its receptor. We filtered this set to include diseases where both the ligand and its receptor were preferentially expressed in distinct cell types.

## ACKNOWLEDGEMENTS

This study was funded by the Israel Science Foundation [317/19 to E.Y.-L].

