## Supplementary Materials for "Affected cell types for hundreds of Mendelian diseases revealed by analysis of human and mouse single-cell data"

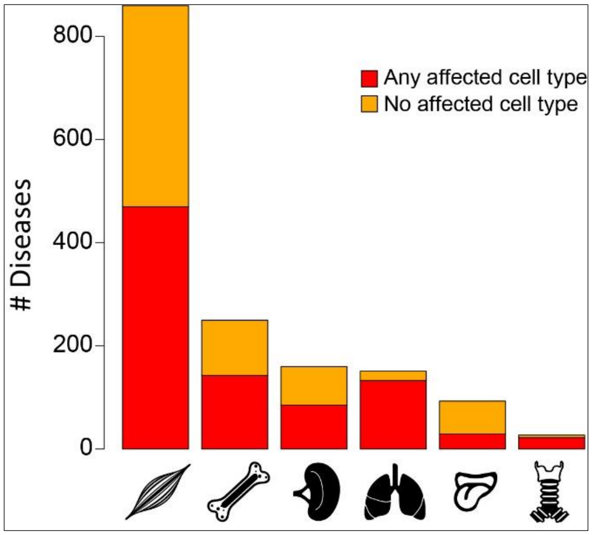


**Figure S1. Disease-affected tissues.** Bar plot of the number of diseases per affected tissue that were either associated with any affected cell type (red) or not (orange).

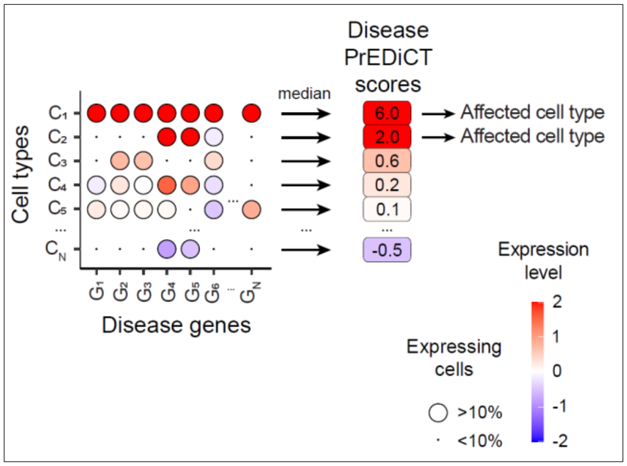


**Figure S2. The PrEDiCT scoring scheme.** To calculate PrEDiCT scores, we first calculated the preferential expression of disease genes in each cell type relative to other cell types of the disease-affected tissue. Next, we set the PrEDiCT score of a disease in each cell type to the median preferential expression of disease genes. Genes expressed (≥0.05 normalized counts) in <10% of the cells per cell type were excluded from calculation. Cell types with PrEDiCT scores ≥2 were considered as affected.

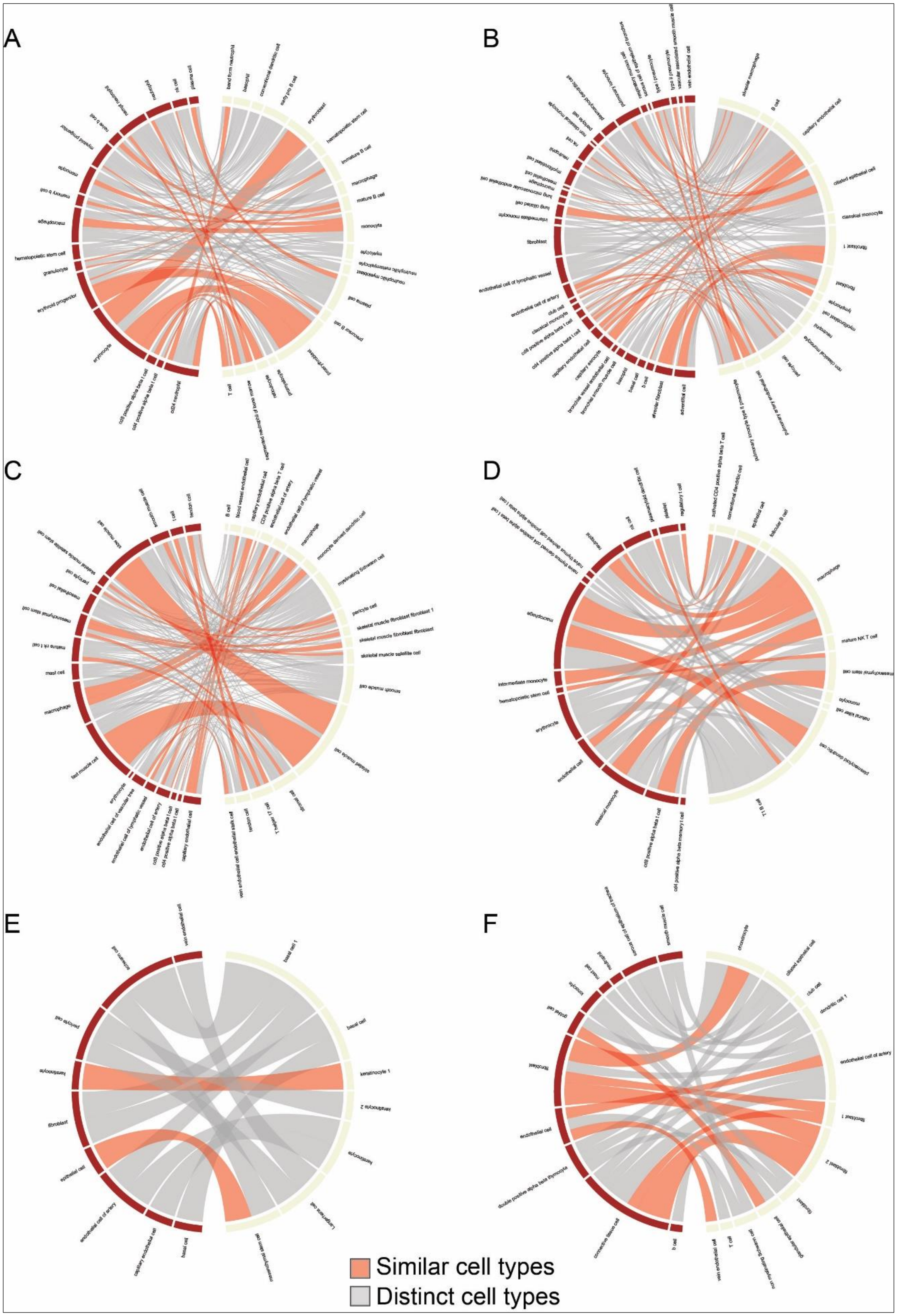
A
B
C
D
E
F
Similar cell types
Distinct cell types

**Figure S3. Affected cell-type similarity between human and mouse tissues.** Circos plots representing similarity between human and mouse cell types (brown and beige, respectively; Table S5). Plots show cell types affected by diseases of the bone marrow, lung, skeletal muscle, spleen, tongue and trachea (A to F, respectively; Table S6). Width and color of lines that connect cell-type pairs between the species indicate the fraction of diseases affecting each cell type and their similarity (red: similar, grey: distinct), respectively.

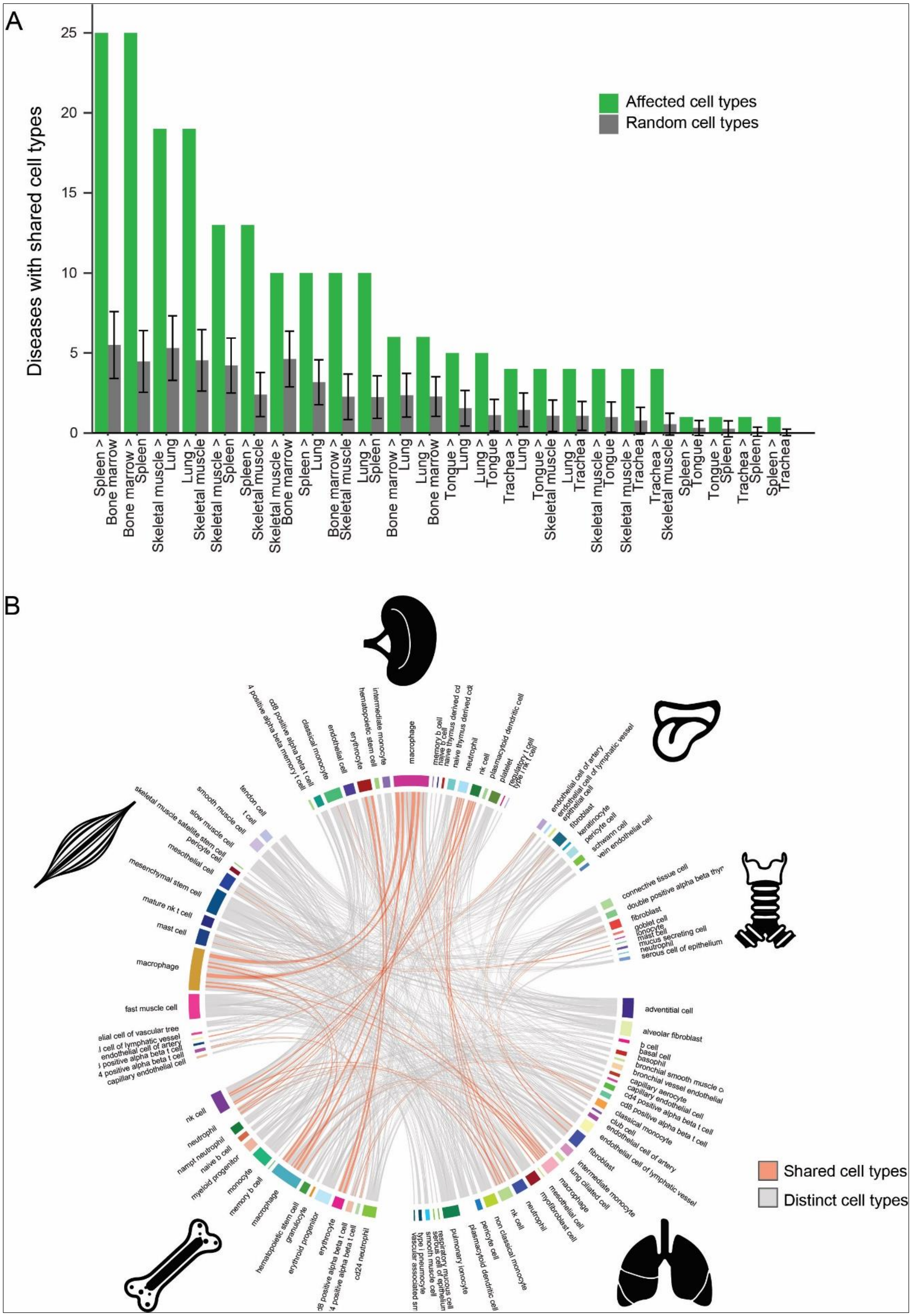
A
Diseases with shared cell types
Affected cell types
Random cell types
B
Shared cell types
Distinct cell types

**Figure S4. Affected cell-type similarity among human tissues.**

A. For each disease, affected cell types in one tissue were shared with another affected tissue (green), more than expected by chance (grey; Permutation test; Methods). Under the bars, tissue (first) and the one it is compared to (second) are shown.

B. Circos plot representing similarity between cell types among human tissues. Width and color of lines that connect cell-type pairs indicate the fraction of diseases affecting each cell type and their similarity (red: similar, grey: distinct; Table S3), respectively.
